# Transmembrane helix 6b links proton- and metal-release pathways to drive conformational change in an Nramp transition metal transporter

**DOI:** 10.1101/792127

**Authors:** Aaron T. Bozzi, Anne L. McCabe, Benjamin C. Barnett, Rachelle Gaudet

**Affiliations:** Department of Molecular and Cellular Biology, Harvard University, 52 Oxford Street, Cambridge, MA 02138, USA

**Keywords:** Nramp, MntH, DMT1, conformational cycle, transition metal transport, cysteine accessibility, proton transport

## Abstract

The natural resistance-associated macrophage protein (Nramp) family encompasses transition metal and proton co-transporters found in organisms from bacteria to humans. Recent structures of *Deinococcus radiodurans* (Dra)Nramp in multiple conformations revealed the intramolecular rearrangements required for alternating access. Here we demonstrate that two parallel cytoplasm-accessible networks of conserved hydrophilic residues in DraNramp—one lining the wide intracellular vestibule for metal release, the other forming a narrow proton-transport pathway—are essential for metal transport. We further show that mutagenic or post-translational modifications of transmembrane helix (TM) 6b, which structurally links these two pathways, impedes normal conformational cycling and metal transport. TM6b contains two highly conserved histidines, H232 and H237. Different mutagenic perturbations for H232, just below the metal-binding site along the proton-exit route, differentially affect DraNramp’s conformational state, suggesting H232 serves as a pivot point for conformational change. In contrast, any tested replacement for H237, lining the metal-exit route, locks the transporter in a transport-inactive outward-closed state. We conclude that these two histidines, and TM6b more broadly, help trigger the bulk rearrangement to the inward-open state upon metal binding and facilitate the return of the empty transporter to an outward-open state upon metal release.

---

Approximately one third of all proteins require specific metal ion co-factors (1), which bind in dedicated sites to stabilize tertiary structures, impart catalytic properties to enzymes, increase protein affinity for other ligands, and enable electron transfer. To maintain metal homeostasis, organisms must acquire metal ions in sufficient quantity from their environment, traffic them to their proper destination, and safely store or excrete any excess to avoid toxicity (2,3). Natural resistance-associated macrophage proteins (Nramps) are transporters that harness the electrochemical energy of proton gradients and the membrane potential to power the uptake of divalent transition metals (4-7). Nramp homologs scavenge manganese in bacteria (8) and acquire and traffic manganese and iron in plants (9) and fungi (10). In mammals, one homolog, Nramp2—also known as Divalent Metal Transport 1 (DMT1)—facilitates dietary iron uptake in the duodenum (11,12) and erythroblast iron-loading in the bone marrow (13,14), while the eponymous homolog, Nramp1, helps phagocytes kill engulfed pathogens, likely by extracting essential metals from phagosomes (15,16).

We developed the *Deinococcus radiodurans* (Dra)Nramp homolog as a model system to understand the general mechanism of this family of transporters (4,17-19). DraNramp is the sole high-affinity Mn^2+^ uptake system (20,21) for a species that maintains an exceptionally high intracellular Mn^2+^ concentration as a resistance mechanism to radiation damage (22), and thus it may be a particularly robust transporter given the likely demand for high expression and activity *in vivo*. Crystal structures of DraNramp in outward-open, inward-occluded, and inward-open states revealed a LeuT-fold—common among secondary transporters (23)—and conformations consistent with an alternating-access model (18,19). Transmembrane segments (TMs) 1, 4, 5, 6, and 10 undergo the greatest displacement relative to each other and to the remaining six TMs, which function as a “scaffold” to support those movements (19,24). Furthermore, while metal transport requires bulk conformational change between outward- and inward-open states, proton uniport occurs through the outward-open—but not inward-open—state (19). Thus, metal ions and protons both transit the external aqueous vestibule to reach their conserved binding site—D56 for protons, and D56 along with N59, M230, and the A53 and A227 carbonyls for metals (19,25)—in the center of the protein, but take separate pathways from there to reach the cytoplasm (19).

Here we identify separate clusters of conserved hydrophilic residues that line each of these pathways. Point mutations to alanine at most of these positions impair DraNramp Mn^2+^ and Co^2+^ transport. Using chemical modifications of a panel of single-cysteine mutants that span TM6, we show that adding steric bulk at many positions along the helix eliminates metal transport, likely by blocking essential conformational rearrangements. Furthermore, alanine replacement of several conserved TM6b residues impedes DraNramp from sampling the outward-open state, as assessed using a single-cysteine reporter. Lastly, using a range of amino acid replacements for two highly conserved histidines, we show that any perturbation of H237 locks the transporter in a transport-inactive outward-closed state, while substitutions of H232 have a range of effects on DraNramp’s transport activity and conformational preferences that suggest H232 may be a pivot point for conformational change. From these results, we conclude that TM6b—and its two histidines in particular—play a critical role in the conformational change process required for metal transport. As the structural connection between the two hydrophilic intracellular substrate-release pathways, TM6b may sense metal binding and/or the ensuant proton transfer to help convert local changes into the bulk conformational rearrangement required to complete the metal transport process.

## Results

### Conserved hydrophilic residues cluster in two networks on DraNramp’s cytoplasmic side

DraNramp consists of 11 TMs, the first 10 of which adopt the canonical LeuT inverted-repeat of two pseudosymmetric five-TM units. We used our previously published alignment of 6878 Nramp sequences that include the canonical “DPGN” and “MPH” motifs (19) and determined the positions at which hydrophilic residues predominate (defined as Ser + Thr + Tyr + Asn + Gln +Asp + Glu + His + Lys + Arg > 80%) (**Fig. S1**). We then mapped these positions onto our outward-open and inward-open DraNramp structures (**Fig. 1**). There is a notable paucity of conserved hydrophilic positions in the external half of the transporter, with only a few hydrophilic residues lining the wide aqueous vestibule that provides access to the metal-binding site (**Fig. 1**). Thus, predominantly hydrophobic packing between TMs 1b, 3, 6a, 8, and 10 seals this external vestibule in the inward-open state (18,19). In contrast, a number of hydrophilic positions flank the conserved metal-binding site in the middle of the membrane, and the cytoplasmic half of the protein is rich in hydrophilic residues. The positions in the protein’s lower half form two extended polar networks. One, between TMs 1a, 2, 5, 6a, and 7, lines the wide intracellular vestibule that serves as the metal-release pathway in the inward-open state (**Fig. 1**)(18). The second, between TMs 3, 4, 8, and 9, forms the narrow pathway that both provides a route for proton uniport in the outward-open state (19) as well as metal-stimulated proton transport (4). These two networks meet at the conserved proton/metal binding site (6,19,25), where they provide parallel exit pathways that facilitate the co-transport of two like-charges (19). The highly conserved TM6b forms the clearest structural connection between these two networks.

**Figure 1.**
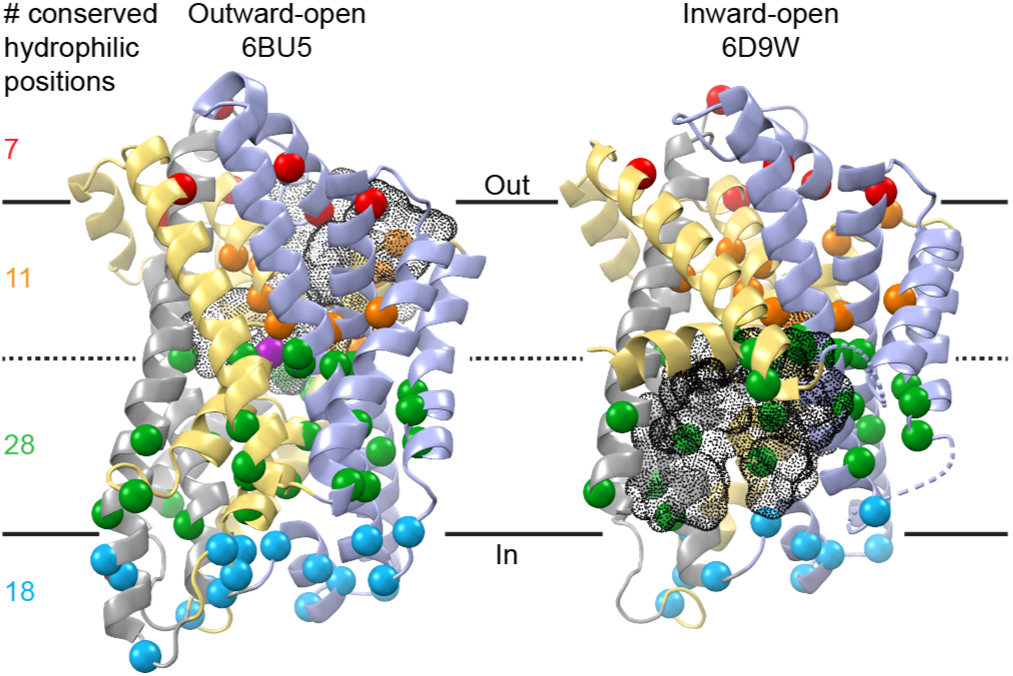
Conserved hydrophilic positions cluster on the intracellular side of DraNramp. Cα positions at which Ser + Thr + Tyr + Asn + Gln +Asp + Glu + His + Lys + Arg > 80% from an alignment of 6878 Nramps are shown as colored spheres on the outward-open Mn^2+^-bound (PDB # 6BU5) and inward-open apo (PDB # 6D9W) structures of DraNramp (19). TMs 1, 5, 6, and 10 are gold, TMs 2, 7, and 11 are gray, and TMs 3, 4, 8, and 9 are blue. The Mn^2+^ substrate is magenta in the outward-open structure. The external and internal vestibules that enable metal entry and release are shown as black mesh. A sequence logo based on the alignment is in Figure S1.

To assess the importance of these residues to the general metal-transport mechanism, we generated a panel of DraNramp point mutants that we expressed in *Escherichia coli* (**Fig. S1** and **S2)**. We then measured relative rates of *in vivo* Co^2+^ transport (WT K_M_ ≈ 1 mM (4)) via our established colorimetric assay (17) and Mn^2+^ transport (WT K_M_ ≈ 3 μM (4,19)) with a new assay in which metal uptake is monitored through the increase in fluorescence of intracellular GCaMP6f (**Fig. S3**). We discuss the results of these experiments below in the context of our recent high-resolution crystal structures of outward-open and inward-occluded DraNramp (19) as we explore the roles of the two polar networks and TM6b to DraNramp function.

### Mutations to non-helical binding-site region impair metal transport

Consistent with other LeuT-fold transporters (26,27), DraNramp uses non-helical regions in the middle of TM1 and TM6 to bind substrates (19,25). In the outward-open state, D56, N59, and the A53 backbone-carbonyl from TM1, and M230 from TM6 coordinate Mn^2+^ (19). In the inward-open state D56, N59, and M230 still coordinate Mn^2+^, but the increasing helicity of TM6a enables the A227 backbone-carbonyl to replace the A53 backbone-carbonyl in the Mn^2+^-coordination sphere in a structure of the related *Staphylococcus capitis* Nramp (25). In addition, Q378 (86% conserved in our alignment, another 11% as N) from TM10 may also directly or indirectly stabilize Mn^2+^ in an occluded conformation (17,19,28). To stabilize the extended unwinding of TM6a in the outward-open state, TM6’s T228 (80% conserved) and TM11’s N426 (99% conserved) donate hydrogen bonds to the unsatisfied backbone carbonyls of I224 and G226 respectively (**Fig. 2A**). In contrast, in the inward-occluded state both sidechains reorient to allow TM6a to extend by two residues, with N426 now donating a hydrogen bond to the carbonyl of C382 to facilitate the toppling of TM10 above P386 (83% conserved) that helps close the external vestibule and allows Q378 to approach the other metal-binding residues (**Fig. 2B**).

**Figure 2.**
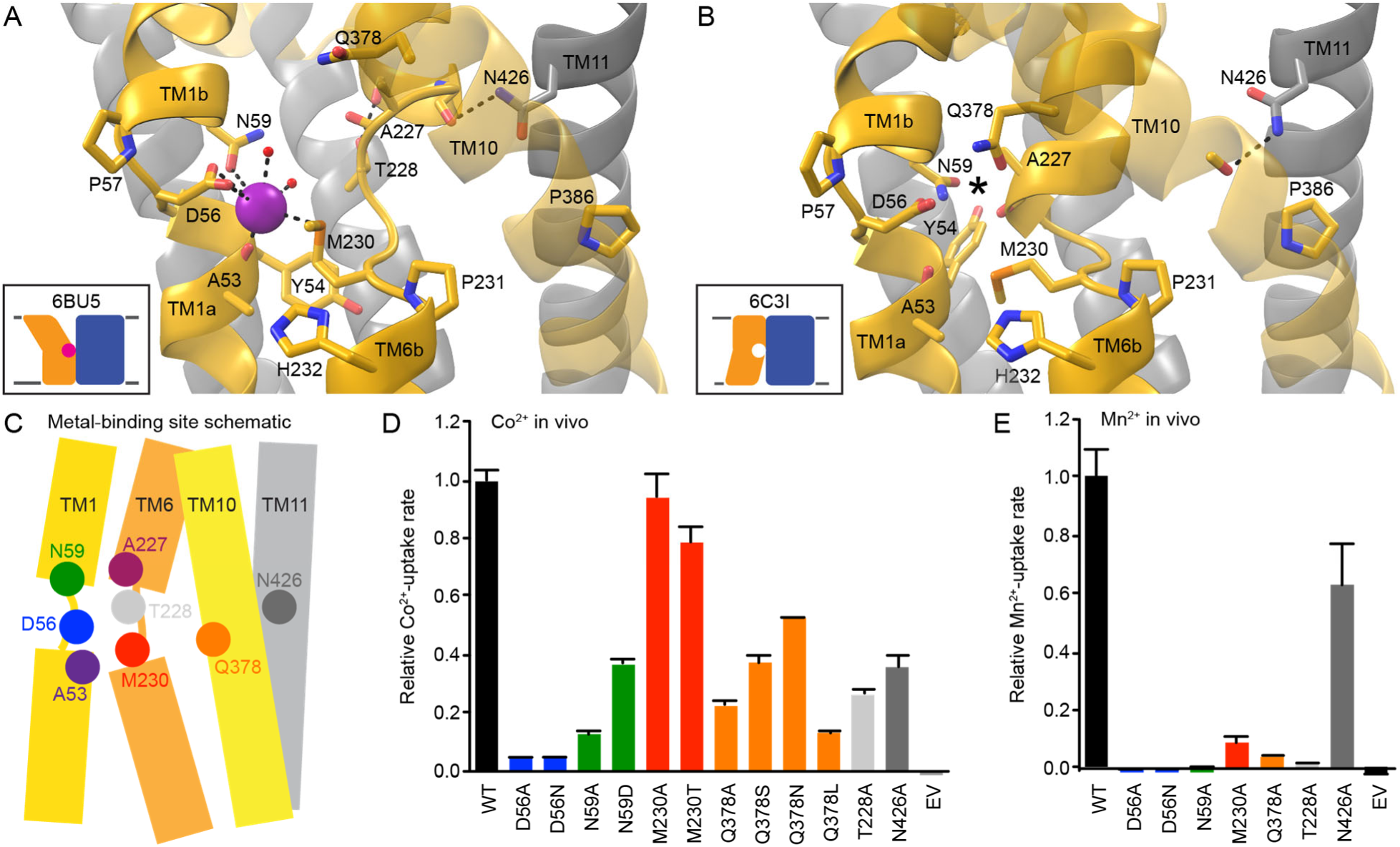
Unwound regions form conserved metal binding site. (A-B) Side view of the metal-binding site region, with the intracellular face of the protein pointing down. For clarity only TMs 1, 2, 6, 7, 10, and 11 are shown. (A) Conserved D56, N59, M230 sidechains along with the A53 carbonyl and two ordered waters coordinate the Mn^2+^ substrate in the outward-open state (PDB # 6BU5). Extended TM6 unwinding is facilitated by the T228 and N426 sidechains donating hydrogen bonds to TM6 backbone carbonyls. (B) In the inward-occluded apo state (PDB # 6C3I), the toppling of TM10 above the conserved P386 enables Q378 to approach the other metal-binding sidechains, while helical extension of TM6a brings the A227 carbonyl into the metal-binding site (* indicates the expected location for a bound metal ion in this and subsequent figures). (C) Schematic of important residues in the unwound metal-binding site. (D-E) Most mutations to conserved binding-site residues impaired relative *in vivo* Co^2+^ (D) and Mn^2+^ (E) uptake rates. Data are averages ± S.E.M. (n ≥ 4 for Co^2+^, n ≥ 6 for Mn^2+^).

As expected, mutations to the conserved residues in the metal-binding site (**Fig. 2C**) were all deleterious to Co^2+^ and Mn^2+^ transport (**Fig. 2D-E**), with the exception of M230 mutants that preserved high Co^2+^ transport as seen previously (4,17,19). Serine and asparagine replacements for Q378 preserved greater activity than alanine or leucine, indicating the importance of a hydrophilic residue at the position. In addition, the mutations T228A and N426A, which remove the hydrogen-bond donating sidechains that support the non-helical metal-binding region, also impaired transport.

### Mutations to inner-gate polar network impair metal transport

The opening of the interior metal-release pathway between TMs 1a, 2, 5, 6b, and 7 proceeds via rearrangements within one of the two networks of highly conserved hydrophilic residues on the cytoplasmic side of the protein (**Figs. 1 and 3E**). In the outward-open structure, Y54 (90% conserved, 10% F) acts as a gate by filling the interface among TMs 1a, 2, and 6b and forms a hydrogen-bonding network that includes Q89 (100% conserved) and H237 (93% conserved) (**Fig. 3A-B**). In the inward-occluded state, Y54 is flipped up away from TM6b towards TM7, anchoring a new hydrogen-bonding network that includes N82 (51% conserved), N275 (100% conserved), and T228 (80% conserved) (**Fig. 3C-D**).

**Figure 3.**
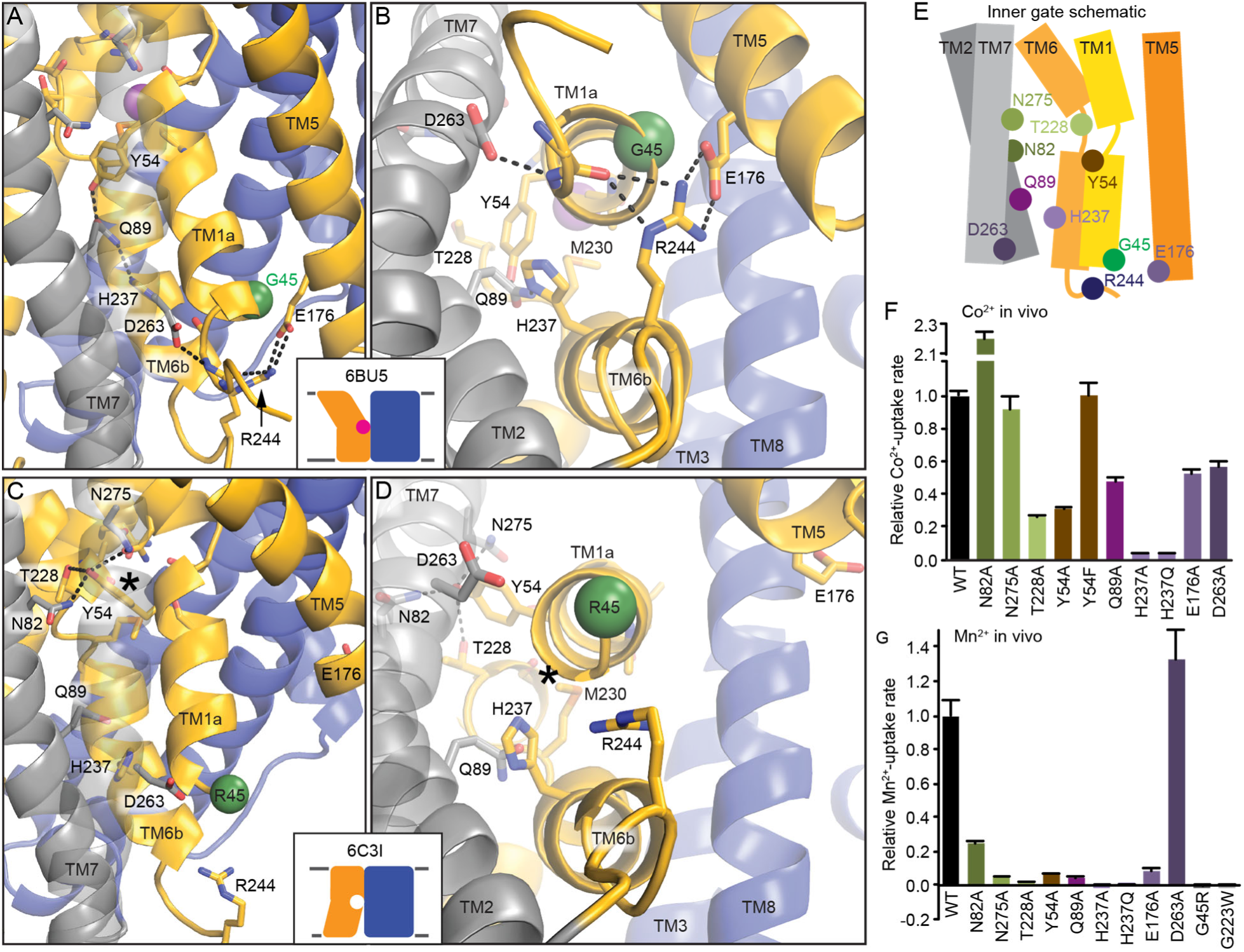
Network of conserved hydrophilic residues rearranges to open intracellular vestibule for metal release. (A-B) Conserved hydrophilic interactions close the inner gate in the outward-open structure viewed from the side (A) or the intracellular face (B). Y54 forms a gate below the bound metal as part of a network that includes Q89 and H237, while lower down R244, E176, and D263 form an extended salt-bridge network that helps lock TM1a in place. (C-D) In the inward-occluded structure viewed from the side (C) or the intracellular face (D), the lower salt-bridge network is disrupted, likely due to the bulky G45R mutation (green sphere shows Cα). Y54 reorients to interact with N82, T228, and N275, and no longer obstructs the metal-binding site from the inside. (E) Schematic of the inner gate network formed by TMs 1a, 2, 5, 6b, and 7. (F-G) Most mutations of the hydrophilic gating residues impaired relative *in vivo* Co^2+^ (F) and Mn^2+^ (G) uptake rates. Data are averages ± S.E.M. (n ≥ 4 for Co^2+^, n ≥ 6 for Mn^2+^).

Further below the metal-binding site, in the outward-open state the E176-R244 (99% and 84% conserved, respectively) salt bridge tethers TM5 to TM6b, while R244 and D263 (75% conserved, 24% E) interact with backbone carbonyls to hold TM1a in place (**Fig. 3A-B**). These inner-gate interactions must be disrupted before TM1a swings to fully open the inward metal-release pathway. Indeed, they are absent in the inward-occluded (and inward-open) structure (**Fig. 3C-D**) with E176 and R244 ending up 20 Å apart. The invariant G45 presses tightly against the E176-R244 pair in the outward-open state, such that any bulkier residue creates steric clashes. Thus the G45R mutation precludes the proper closing of the inner gate and prevents sampling of the outward-open state (18,19), explaining the loss-of-function phenotype (18) that causes anemia in humans with the analogous mutation in DMT1 (29).

Mutations to these conserved hydrophilic residues involved in opening and closing the inner gate were mostly deleterious to metal transport (**Fig. 3F-G**). However, N275A and Y54F did not significantly reduce, and N82A actually enhanced, the rate of Co^2+^ transport, while D263A did not impair Mn^2+^ uptake.

### Mutations to proton-transport pathway polar network impair metal transport

Proton transport occurs via a pathway separate from the intracellular metal-release route, which remains closed to bulk solvent in proton-transporting, outward-locked mutants (19). On the opposite side of the metal-binding site from Y54, which forms the first barrier to metal release, begins a network of highly conserved hydrophilic residues. This network includes at least seven potentially protonatable sidechains and leads from proton- and metal-binding D56 (19) through a tight corridor between TMs 3, 4, 8, and 9 to the cytoplasm (**Fig. 4A-B**) to provide a route for proton transport (4,19). In contrast to the external and intracellular vestibules proposed as metal entrance and release pathways, the helices and residues within this polar network, with the exception of the cytoplasmic end of TM4, undergo little rearrangement between our outward-open, inward-occluded, and inward-open structures (18,19).

**Figure 4.**
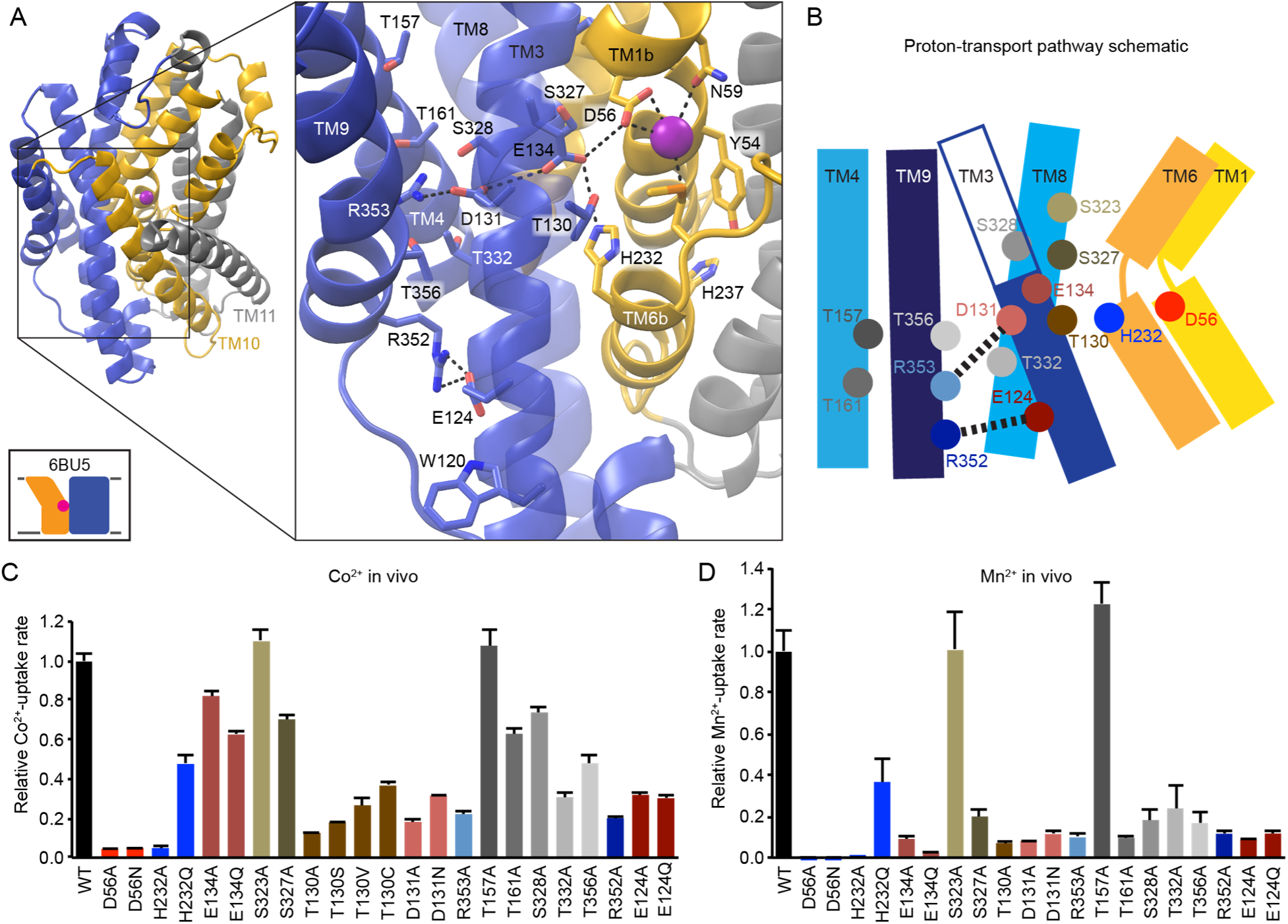
Proton-transport pathway leads through conserved polar network from metal-binding site to cytoplasm. (A) View from an angle above the membrane looking down into the extracellular vestibule in the outward-open structure, with zoomed-in inset (with TM10 and TM11 omitted for clarity), and (B) schematic showing a network of conserved hydrophilic residues leading from the metal-binding site into the cleft between TMs 3, 4, 8, and 9. H232 and E134 abut metal-binding M230 and D56, providing a connection to the D131-R353 and R352-E124 salt-bridge pairs. Many moderately conserved hydrophilic residues line the surrounding passageway. (C-D) Mutations to many residues in the proton-transport pathway impaired relative *in vivo* Co^2+^ (C) and Mn^2+^ (D) uptake rates. All data are averages ± S.E.M. (n ≥ 4 for Co^2+^, n ≥ 6 for Mn^2+^).

Highly-conserved residues surrounding the metal-binding D56 includes H232 on TM6b (100% conserved) and E134 (TM3, 98% conserved)— which are essential to Nramp proton uniport and proton-metal co-transport (4,6,19,28)—along with T130 (TM3, 69% conserved) and S327 (TM8, 92% conserved). Across from E134 lies a conserved salt-bridge pair: D131 (TM3, 93% conserved) and R353 (TM9, 78% conserved), with D131, which is required for proton transport, the likely proton-transfer point after incoming metal causes D56 to deprotonate (4,19). Approximately 9 Å below, a second conserved salt bridge, E124-R352 (94% and 87% conserved, respectively), links the same two helices, with several moderately conserved serines and threonines from TMs 4, 8, and 9 in between. Hydrophobic residues around the salt-bridge network and below E124 may help restrict the accessibility of bulk solvent; in previous work we detected a slight accessibility along that face of TM3 only up to E124 (18).

Alanine replacement at most positions within this extended polar network generally impaired Co^2+^ and Mn^2+^ transport (**Fig. 4C-D**). Interestingly, alanine replacement of E134—essential to voltage dependence and proton-metal coupling (4)— preserved significant Co^2+^ uptake, while H232A eliminated transport. T130, which flanks the interface of the crucial D56-E134-H232-M230 tetrad along the proton-transfer route to D131, is particularly important for metal transport, with larger replacements such as T130C preserving greater activity than T130A, indicating that steric bulk here likely aids optimal binding-site alignment. Mutations to S327 and S328 (20% conserved, 74% T), which also line the proton-transfer route and may be a remnant of an ancestral Na^+^-binding site (19,30), impaired Mn^2+^ uptake but were less deleterious to Co^2+^ transport. Mutations that disrupt the E124-R352 and D131-R353 salt-bridge pairs greatly reduced transport of both substrates. For T157 (85% conserved), T161 (35% conserved), T332 (83% conserved), and T356 (94% conserved), which cluster between the two TM3-TM9 salt bridges, alanine substitution also impaired transport, but to a lower degree than mutations to the charged residues. Overall, the intact TM3, TM4, TM8, TM9 polar network that provides the proton exit pathway was essential to efficient DraNramp metal transport.

### TM6b drives the conformational change required for metal transport

TM6b forms the main structural connection between the two parallel polar networks that diverge from the DraNramp binding site into otherwise distinct structural elements of the protein (**Fig. 1**). In the outward-open and inward-occluded states, TM6b is closely intertwined with TM1a (19), while this interaction is disrupted in the fully inward-open structure (18). Adding steric bulk to positions on TM1a that would prevent the observed tight packing with TM6b eliminated metal transport and locked the transporter in an outward-closed conformation (18). In addition, mutations at several positions on TM6 impaired high-affinity Mn^2+^ transport in *E. coli* Nramp (31). Based on these previous findings, we postulated that TM6b might play an essential role in corralling TM1a to fully close the metal pathway’s inner gate as a prerequisite for opening the external vestibule.

To test this hypothesis, we measured the Co^2+^ transport ability of a panel of single-cysteine mutants spanning TM6 (**Fig. 5A**). Introducing cysteines along TM6a (A217 to A227) did not greatly affect function, except at the invariant G226. However, for ten consecutive positions from the unwound region at T228 through the first half of TM6b to H237, cysteine substitution moderately or severely impaired metal transport, particularly at the highly conserved L236 (93% conserved) and H237 (93% conserved). In contrast, cysteine substitution in the second half of TM6b (S238 to Q242) did not notably diminish transport, except at L240.

**Figure 5.**
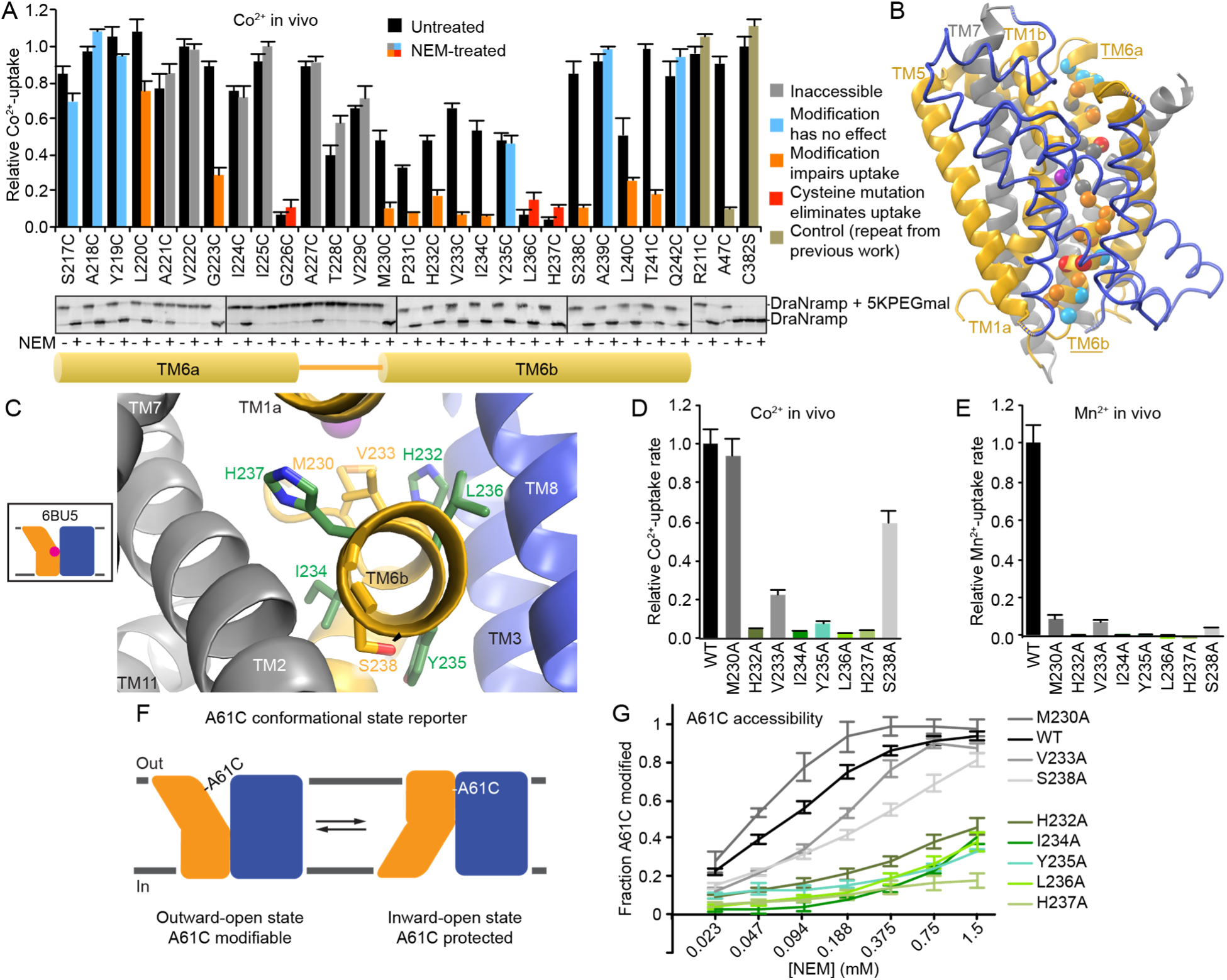
TM6 plays instrumental role in DraNramp bulk conformational change. (A) Relative *in vivo* Co^2+^ uptake of single-cysteine mutants (which also contain the C382S mutation to remove the lone endogenous cysteine) along TM6 (black bars) showed functional importance of TM6b. Cysteine modification with NEM (colored bars) further reduced transport for most positions on TM6b, likely by impeding essential conformational rearrangements. Data are averages ± S.E.M. (n = 3). Western blots indicate which cysteines could be fully modified: preincubation with NEM prevents the upward shift caused by subsequent 5K-PEG maleimide modification under denaturing conditions. (B) Results from panel A mapped onto the outward-open structure to illustrate where NEM modification is tolerated or not. (C) Conserved sidechains along TM6b viewed from the cytoplasm. (D-E) Alanine scanning of TM6b residues reduced or eliminated relative *in vivo* Co^2+^ (D) and Mn^2+^ (E) uptake. Data are averages ± S.E.M. (n ≥ 4 for Co^2+^, n ≥ 6 for Mn^2+^). (F) A61C acts as a reporter for sampling of the outward-open state, as this position is fully protected from NEM modification in the inward-open state. (G) Many TM6b alanine mutants protected A61C from NEM modification, indicating an impairment of conformational cycling. Data are averages ± S.E.M. (n ≥ 4).

Next, we investigated the effects of bulky NEM modification along TM6 (**Fig. 5A-B**). Briefly, we used a two-step labeling protocol, in which NEM was applied to *E. coli* expressing the single-cysteine DraNramp construct. Cells were then lysed and protein denatured before adding 5K-PEG maleimide to modify previously unlabeled cysteines and cause a gel shift on Western blots. Most positions on TM6a were not full modified, indicating a highly constricted environment. In contrast, all positions from M230 and below were completely NEM-labeled, likely from the protein sampling a conformational state with a large cytoplasmic aqueous vestibule as seen in our original inward-open structure (18). Adding the bulky NEM moiety along the external vestibule at L220 or G223 impaired metal transport, likely by impeding the proper closing of that vestibule, consistent with our previous results for tryptophan substitution at those positions (19). Interestingly, NEM modification further reduced or eliminated Co^2+^ transport for eight positions on TM6b (M230-I234, S238, L240, T241), but not at Y235, A239, or Q242 (**Fig. 5A-B**). Those latter positions face the “scaffold” domain (TM3 and TM8). These results suggest that the lower part of TM6b (Y235 and below) does not snugly interact with TM3 and TM8—which, along with TM4 and TM9 form the pathway for proton transport (19)—in any essential conformation. However, on all other sides of TM6b, which face TMs 1a, 2, 7, and 10, NEM modification eliminated any residual Co^2+^ transport, likely because the adduct created steric clashes that block essential conformational rearrangements, such as inward movement of TM1a. Thus, while TM6b itself does not reorient dramatically in our prior crystal structures, it may form a nexus for the rearrangements of other essential moving parts in the DraNramp transport cycle, or else move significantly itself in an as-yet-uncaptured conformational state.

To further understand the role of TM6b, we measured Co^2+^ and Mn^2+^ transport for a panel of alanine substitutions from M230-S238 (**Fig. 5C-E**). For this panel we also measured A61C accessibility to assess the mutants’ conformational cycling ability (**Fig. 5F-G**)—this single cysteine reporter on TM1b is only accessible to NEM modification in DraNramp’s outward-open conformation (18,19). Most mutations removing bulky sidechains—H232A, I234A, Y235A, L236A, and H237A—reduced sampling of the outward-open state and impaired or eliminated metal transport. In contrast with our prior observations regarding TM1a (18), not only adding bulk but also removing bulk from TM6b prevented the conformational change needed for metal transport, underscoring the role that its mix of hydrophilic and hydrophobic residues likely play in stabilizing helix packing. The native residues of TM6b may therefore assist in the closing of the intracellular vestibule, which is likely a prerequisite process for opening of the external vestibule to reach the outward-open state.

### TM6b histidines link substrate release pathways to drive conformational change

The clearest structural connection between the two polar networks that form the adjacent intracellular metal- and proton-release pathways is TM6b, on which H232 and H237 occupy opposite faces of a highly conserved helix below the metal-binding site (**Fig. 5C and 6A**). Others have argued that one or both of H232 and H237 have roles in metal binding, proton transport, and/or pH regulation in various Nramp homologs (6,32-34). H232’s position at the protein’s core renders it unsuitable for direct protonation as the congested environment disfavors a net charge on the imidazole (19,28). This residue is nevertheless essential to high-affinity Mn^2+^ transport, H^+^ uniport, and proton-metal co-transport in DraNramp (4,19) and *Eremococcus coleocola* Nramp (6). We previously proposed it plays a key role, along with the adjacent E134, in stabilizing a proton transfer from D56 to D131 (4). H237, located two helical turns more intracellular on TM6b from the metal-binding M230, is quite distant from the metal-binding site (13.4 Å from the bound Mn^2+^ in the outward-facing state) on the TM6b face most distant from the proton-release pathway.

To further probe the importance of these two residues, we tested Co^2+^ transport and A61C accessibility for a variety of sidechain replacements at H232 and H237 (**Fig. 6B-C**). All tested H237 substitutions yielded inward-locked transporters that did not transport metal (**Fig. 6B-C**), illustrating that residue’s indispensable role in stabilizing the outward-open state. An alanine substitution of the invariant Q89—the H237 hydrogen-bond partner in the outward-open state—had similar but less severe effects.

**Figure 6.**
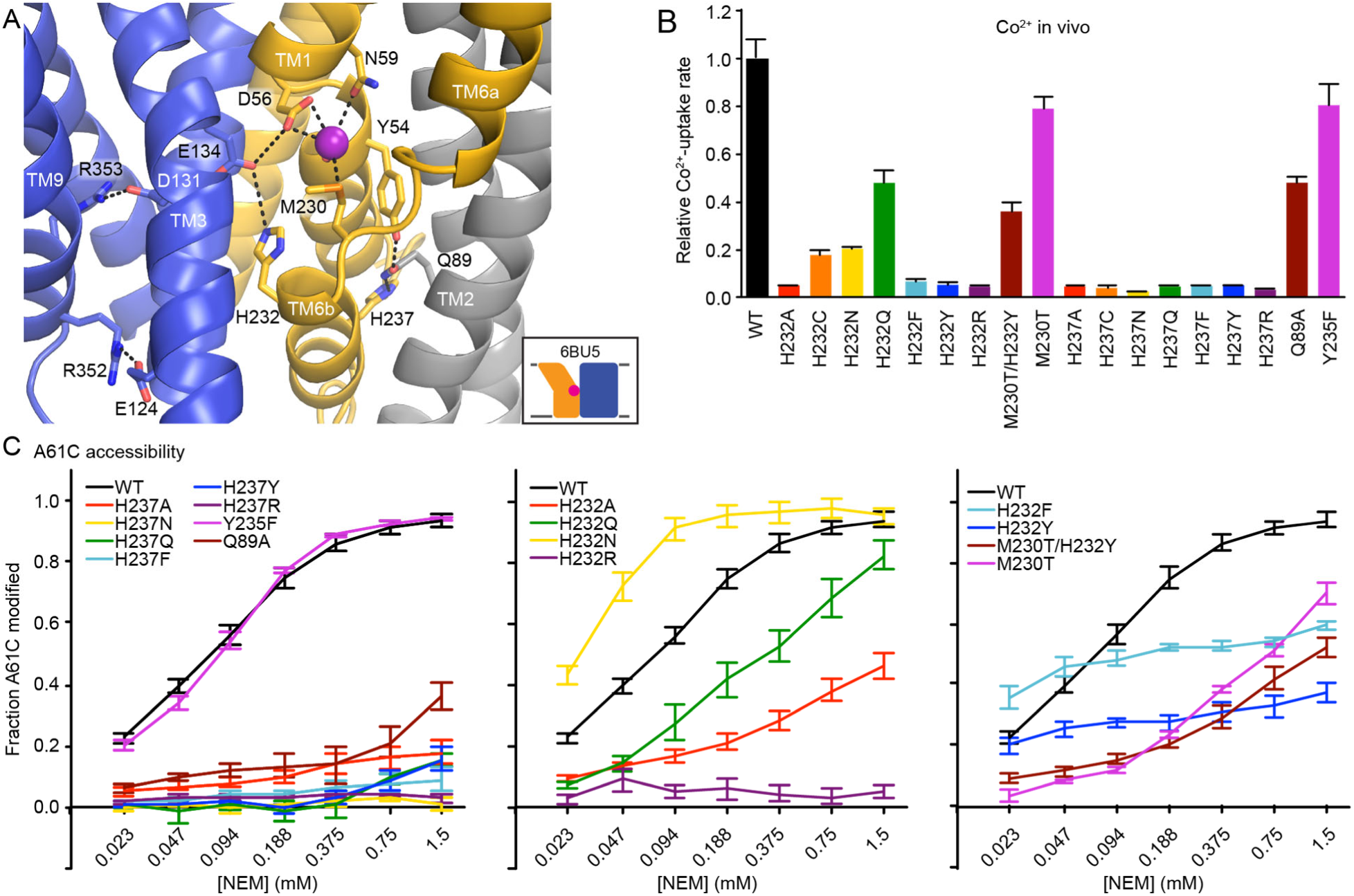
TM6 histidines govern DraNramp conformational state. View from the periplasm of outward-open DraNramp highlighting the key positions of H232 and H237 in relation to the metal-binding site and proton-transport pathway. For clarity TMs 10 and 11 are removed.(B) All tested mutations to H232 impaired relative *in vivo* Co^2+^ uptake rates, with H232Q the only significantly active mutant. All tested mutations to H237 eliminated relative *in vivo* Co^2+^ uptake. (C) NEM accessibility experiments showed all mutations to H237 rendered A61C fully protected, indicating an outward-closed state. (G-H) Mutations to H232 perturbed A61C accessibility, indicating disparate effects on the conformational equilibrium. All data are averages ± S.E.M. (n ≥ 4).

For H232 the results varied, although all tested mutants besides H232Q profoundly impaired metal transport (**Fig. 6B**). H232N and H232Q slightly increased and decreased A61C accessibility, respectively, while H232R locked the transporter in an outward-closed state (**Fig. 6C**). Surprisingly, NEM labeling at A61C of H232F and H232Y matched or exceeded WT at low NEM concentrations, but then plateaued at only ∼50% and ∼25% respectively, whereas A61C labeling of WT reaches completion (**Fig. 6C**). One interpretation of this result is that the H232F and H232Y variants were trapped in a mixture of inward- and outward-open states, with an energy barrier too high for rapid interconversion, thus explaining the lack of metal transport. Interestingly, the M230T/H232Y double substitution, which changes the TM6 “MPH” motif to the “TPY” found in Nramp-related Mg^2+^- and Al^3+^-transporters (35,36), may relieve this jam, as it partially restored both A61C labeling and metal transport (**Fig. 6B-C**), emphasizing the structural and functional connection between H232’s position and the metal-binding site. This conserved histidine lies at the interface of the mobile and scaffold regions of the protein along the proton pathway, and thus could serve as a pivot point for bulk conformational change upon sensing metal binding or proton transfer.

## Discussion

Previously, we demonstrated the importance of an alternating-access mechanism to Nramp metal transport (18,19) and elucidated the role of a conserved salt-bridge network in enabling proton uniport and proton-metal co-transport (4). Here we illustrated the importance of two parallel cytoplasmic networks of conserved hydrophilic residues to *in vivo* DraNramp metal transport (**Fig. 7**). One polar network extends below the metal-binding site to form an inner gate that rearranges to allow metal release in the inward-open state (**Fig. 3**). A second polar network encapsulates the essential salt-bridge network and provides a route for proton exit to the cytoplasm (**Fig. 4**). The highly conserved TM6b, which forms the structural link between these two networks, is essential to the conformational change process (**Fig. 5**). TM6b’s two conserved histidines, H232 and H237, are particularly important in the control of the transporter’s conformational state (**Fig. 6**) and likely are instrumental in driving the switch to the inward-open state upon metal-binding (H232) and promoting the return of the empty carrier to the outward-open state after metal release (H237).

**Figure 7.**
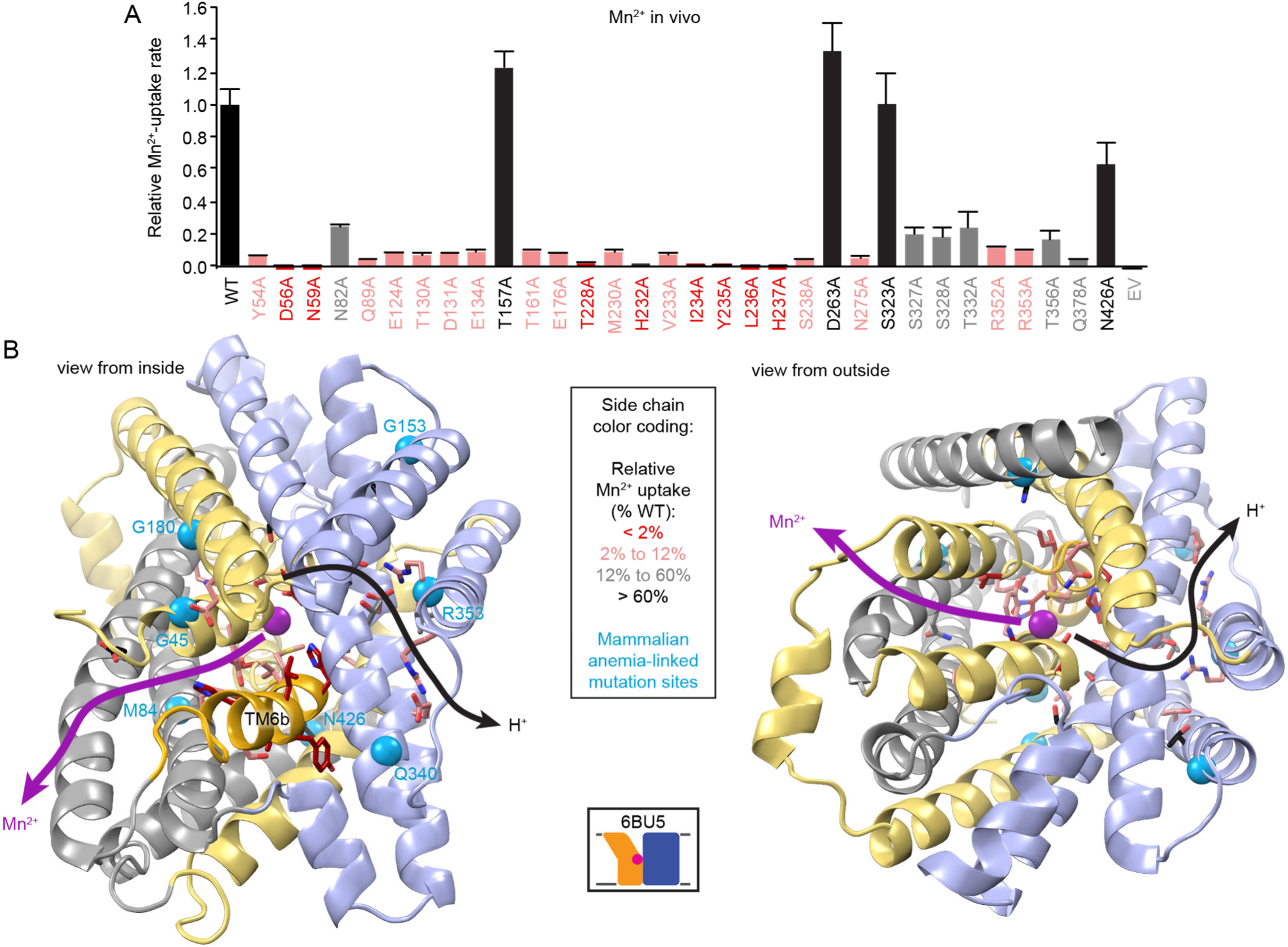
TM6b links two conserved polar networks. (A) Mn^2+^ uptake data for a DraNramp alanine mutant panel. Data are averages ± S.E.M. (n ≥ 6), repeated from Figures 2-5. (B) DraNramp outward-open structure showing sidechains of tested residues as sticks color-coded by the severity of Mn^2+^-transport impairment caused by mutation to alanine. TM6b is highlighted in deeper yellow. Seven mammalian anemia-causing mutations are indicated as cyan spheres. The proposed separate pathways for Mn^2+^ and H^+^ transport to the cytoplasm are shown as arrows.

The inherent asymmetry of Nramps—a hydrophobic outer gate and two hydrophilic cytoplasmic networks, one forming the inner gate for metal release, the second forming a parallel proton-transport pathway—may be an evolutionary adaptation to promote outside-to-inside transport. The hydrophilic inner gate residues may position water molecules to facilitate eventual metal re-hydration and release upon conformational change, and we observed just such an ordered water tethered by H232 in our outward-open structure (**Fig. 6A**) (19). In contrast, the hydrophobic packing above the metal-binding site should prevent any similar assistance in the reverse direction, thus guarding against metal efflux. In addition, there is no structural equivalent to the parallel proton-transport network in the external half of the protein, which provides for the spatial separation of the two like-charge co-substrates during the transport process from outside-to-inside (4,19).

Our functional data in **Figure 5** indicate an essential role for TM6b in conformational change and metal transport not yet captured by crystallography. Superpositions of the available DraNramp crystal structures did not reveal significant displacement of TM6b in the conformational change from outward-open to inward-occluded (Cα RMSD of 0.7 Å) (19). While a slightly-larger difference was seen from inward-occluded to inward-open (Cα RMSD of 2.1 Å) the electron density for the lower half of TM6b (below L236) was not well-resolved in the inward-open structure (19). Analogous structural comparisons of the inward-open *S. capitis* Nramp (34% identical / 56% similar to DraNramp) and the outward-open *E. coleocola* Nramp (33% / 52% to DraNramp, 47% / 64% to *S. capitis* Nramp) also indicated little displacement of TM6b (6). As bulky modification on TM6b facing TMs 1a, 2, 7, and 10 (**Fig. 5A-B**) eliminated metal transport, perhaps those helices reorient significantly relative to TM6b. However, both structural superpositions and distance-difference matrix calculations revealed little relative displacement between TMs 6b and TM2, 7, or 10 (19), which *S. capitis* and *E. coleocola* Nramp comparisons also corroborated (6). In contrast, our analyses did highlight the ∼45° displacement of TM1a in the inward-open state compared to its position in the inward-occluded and outward-open states (19). As would be expected, adding steric bulk on TM1a eliminated metal transport and trapped the protein in an outward-closed conformation, thus validating the functional importance of TM1a’s movement (18). The similar functional results with TM6b therefore suggest a significant displacement for that helix at some point in the transport cycle.

Previous work with the distantly related structural homolog LeuT, a Na^+^-driven amino acid transporter, supports the likelihood of an essential TM6b motion not apparent from crystal structures. Comparisons of LeuT in inward- and outward-open states showed little displacement of TM6b or the neighboring parts of TMs 2, 7, and 10 (6,37). However, double electron-electron resonance (DEER) measurements with LeuT indicated a significant motion for TM6b (and TM7) between conformational states, likely as part of opening and closing the inner gate (38). In addition, molecular dynamics (MD) simulations with LeuT showed more substantial rearrangements for TM6b during substrate release (39,40). Cysteine-modification experiments (41) and MD simulations (42) with LeuT’s close homologs corroborate TM6b’s importance to conformational change. In addition, mutation to alanine of a highly conserved tyrosine on TM6b, which disrupts a hydrophilic inner gate network, caused the protein to relax into the inward-open state, as captured in the crystal structure (37) and confirmed with fluorescence resonance energy transfer (FRET) (43) and DEER data (38). The many apparent similarities between TM6b in DraNramp and LeuT may reflect an already-evolved functional role for this helix in the ancestral precursor to both transporters, which was then preserved even through the divergence of other conformational rearrangements as well as their substrate type and coupling mechanism.

Nramp crystal structures of *S. capitis* and *E. coleocola* homologs revealed analogous polar networks to those seen in DraNramp (6,25). In addition, previous functional studies with the *E. coli* homolog (51% identical and 72% similar to DraNramp) confirmed the importance of the conserved polar networks and TM6b to the transport mechanism. Two studies (31,44) showed deleterious effects for mutations to D34 (D56 in DraNramp), P35 (P57), G36 (G58), N37 (N59), G205 (G226), A206 (A227), and M209 (M230) within the metal-binding site region; E154 (E176) and D238 (D263) within the inner gate network; E102 (E124), D109 (D131), and E112 (E134) in the proton-transport pathway; and P210 (P231), H211 (H232), L215 (L236), H216 (H237), and S217 (S238) on TM6b—which the authors also proposed as a likely key player in the conformational change process (31). Two additional studies (32,45) showed that mutations to the aforementioned D34, N37, H211, and H216 impaired metal uptake, as did mutations to N250 (N275) in the inner gate network and N401 (N426) in the unwound region. Additionally, many of these conserved residues were also shown to be important to metal transport in *Arabidopsis thaliana* Nramp3 (46).

Interestingly, mammalian anemia-causing DMT1 mutations cluster within the two identified polar networks (**Fig. 7B**). The human DMT1 (26% identical, 48 % similar to DraNramp) mutant N491S (N426 in DraNramp) (47) truncates the asparagine that differentially stabilizes the non-helical metal-binding site region (**Fig. 2**). The ΔV114 (M84) (48) mutation shifts the registry of TM2 within the protein’s metal-release pathway. The mutants G75R (G45) on TM1a (29) and G212V (G180) on TM5 (48) in human DMT1 add steric bulk at positions abutting the highly-conserved E176-R244 salt bridge, which forms part of the inner gate that closes the metal-release pathway in the outward-open state (**Fig. 3**). In addition, the human DMT1 mutant R416C (R353) on TM9 (49,50) removes a highly-conserved charged residue from the proton-transport pathway, while the mouse and rat DMT1 mutant G185R (G153) on TM4 (12,51) adds an additional positive charge in the vicinity of that network (**Fig. 4**). The human DMT1 mutant E399D (Q340) on TM8 (48), which makes a conservative substitution in the proton-transport pathway polar network, preserved metal transport function (52).

In addition, designed mammalian DMT1 mutants to D86 (D56, TM1) (17,25,33), N89 (N59, TM1) (25), M265 (M230, TM6) (17,25,28), and N443 (Q378) (28) in the metal-binding site disrupted metal transport, as did mutations to E154 (E124, TM3) (33), D161 (D131, TM3) (33), E164 (E134, TM3) (28), and R416 (R353, TM9) (33,49) in the proton-transport pathway. The TM6b histidines H267 (H232) and H272 (H237) were the focus of two prior studies (33,34). One showed that lowering the external pH rescued some Fe^2+^ transport activity for H267A/C and H272A/C mutants, but not for H267R and H272R replacements, which lacked any activity (33). The authors proposed that these residues directly protonate/deprotonate to regulate transport and conformational change. An alternative explanation consistent with our DraNramp findings is that while the bulky arginine replacements trap the transporter in an inward-locked conformation that prevents transport, the smaller replacements simply shift the conformational equilibrium to disfavor the outward-open state. If a protonation event perhaps stabilizes human DMT1’s outward-open state analogously to how Na^+^ binding stabilizes LeuT’s outward-open conformation (53,54), then lowering the external pH might compensate some for the H267A/C and H272A/C mutants, thus explaining the partial functional recovery. The second study showed a similar reduction of Fe^2+^ transport for H267A/N/D and H272R mutants, but obtained more complicated results with H272A (34). The latter mutant altered relative metal preferences to favor Zn^2+^, eliminated ΔpH stimulation of Fe^2+^ transport, increased the rate of proton uniport, and decoupled co-transport such that adding Fe^2+^ inhibited rather than stimulated H^+^ influx (34). The broad effects of H272A in mammalian DMT1 echo the results of perturbing the TM3-TM9 salt-bridge network, which alter the voltage and ΔpH dependence of metal transport rate as well as proton-metal co-transport stoichiometry in DraNramp (4,19), suggesting that this residue may have an additional essential mechanistic role beyond closing the inner gate to stabilize the outward-open state, at least in some Nramp homologs.

The two identified cytoplasmic polar networks that provide the pathways for metal release and proton transport, and the highly conserved TM6b that connects them, are thus clearly essential to the proper function of our model system DraNramp, as well as *E. coli* Nramp and mammalian DMT1. In particular, our results here, in combination with previously published structural and biochemical data, clearly establish that the integrity of TM6b is crucial to the proper conformational cycling required for efficient import of transition metal into the cytosol by Nramp transporters.

## Experimental procedures

### Cloning of DraNramp

WT and mutant DraNramps were cloned in pET21-N8H as described (18). For the manganese uptake assay, the fluorescent Ca^2+^-sensor GCaMP6f (55) was inserted into pETDuet in the first multiple cloning site using NcoI and NotI cut sites. WT DraNramp and point mutants were inserted into the second multiple cloning site of pETDuet using NdeI and XhoI cut sites, with the vector modified to insert an N-terminal 8xHis-tag that ends in the NdeI site. Mutations were made using the Quikchange mutagenesis protocol (Stratagene) and were confirmed by DNA sequencing. Single-cysteine constructs also included the C382S mutation to remove the lone endogenous cysteine. The C41(DE3) *E. coli* strain was used for protein expression and *in vivo* assays.

### In vivo metal transport assays

Cobalt uptake assays in *E. coli* were performed as described (17,18). For cysteine pre-modification experiments, cells were plated, incubated in labeling buffer (100 mM Tris pH 7.0, 60 mM NaCl, 10 mM KCl, 0.5 mM MgCl_2,_ 0.75 mM CaCl_2_) with or without 3 mM NEM for 15 min at RT, quenched with 10 μL of 200 mM L-cysteine, then washed twice in assay buffer before assaying metal uptake. To measure relative rates of manganese uptake, DraNramp and the fluorescent Ca^2+^-sensor GCaMP6f (55) were co-expressed. The metal-binding EF-hand domain of calmodulin that comprises the Ca^2+^-sensor in GCaMP6f also binds Mn^2+^ in the same two binding sites as Ca^2+^, with a slightly different coordination geometry, and at an approximately 2-fold lower affinity (K_D_ = 13 μM)(56). For this assay, 15 ml of lysogeny broth + 100 μg/mL ampicillin was seeded 1/50 from overnight cultures. Cells were grown at 37°C until OD ≈ 0.6, then induced with 100 μM IPTG and 150 μM EDTA (to sequester trace Mn^2+^ in the media), with protein expression continuing for 2.5 hr at 37°C. Cells were pelleted and washed twice with assay buffer (50 mM HEPES pH 7.3, 60 mM NaCl, 10 mM KCl, 0.5 mM MgCl_2_, 0.217% glucose), and resuspended at OD = 5.26 in assay buffer. Cells were plated 190 μL/well into black clear-bottom plates (Greiner). Baseline fluorescence (λ_ex_ = 470 nm; λ_em_ = 520 nm) was measured on a FlexStation3 (Molecular Devices) for 1 min, before 10 μL 2 mM MnCl_2_ was added (100 μM final concentration) and fluorescence was measured for another 4 min at room temperature (**Fig. S3**). To determine the maximum fluorescence signal, 10 mM CaCl_2_ was added to separate 190 μL aliquots of each mutant and the end-point fluorescence was recorded after a 7-minute incubation. To calculate relative manganese uptake, the pre-metal addition baseline average fluorescence was subtracted from all timepoints, with the residual fluorescence then divided by the corresponding maximum fluorescence increase in the presence of Ca^2+^. Average initial rates were normalized to the WT.

### Cysteine accessibility measurements

Cysteine accessibility measurements were performed as described (18,19).

## Acknowledgements

We would like to thank Wilhelm Weihofen, Lukas Bane, Jack Nicoludis and other Gaudet lab members for helpful discussions. This work was funded in part by NIH grant 1R01GM120996 (to R.G.).

## Conflict of interest

The authors declare that they have no conflicts of interests with the contents of this article. The content is solely the responsibility of the authors and does not necessarily represent the official views of the National Institutes of Health.

## Author Contributions

R.G. oversaw and designed the research with A.T.B. and A.L.M.; A.L.M. and B.C.B created reagents and performed preliminary experiments; A.T.B. performed all functional assays and analyzed the resulting data. A.T.B. and R.G. wrote the manuscript.

---

**Figure S1.**
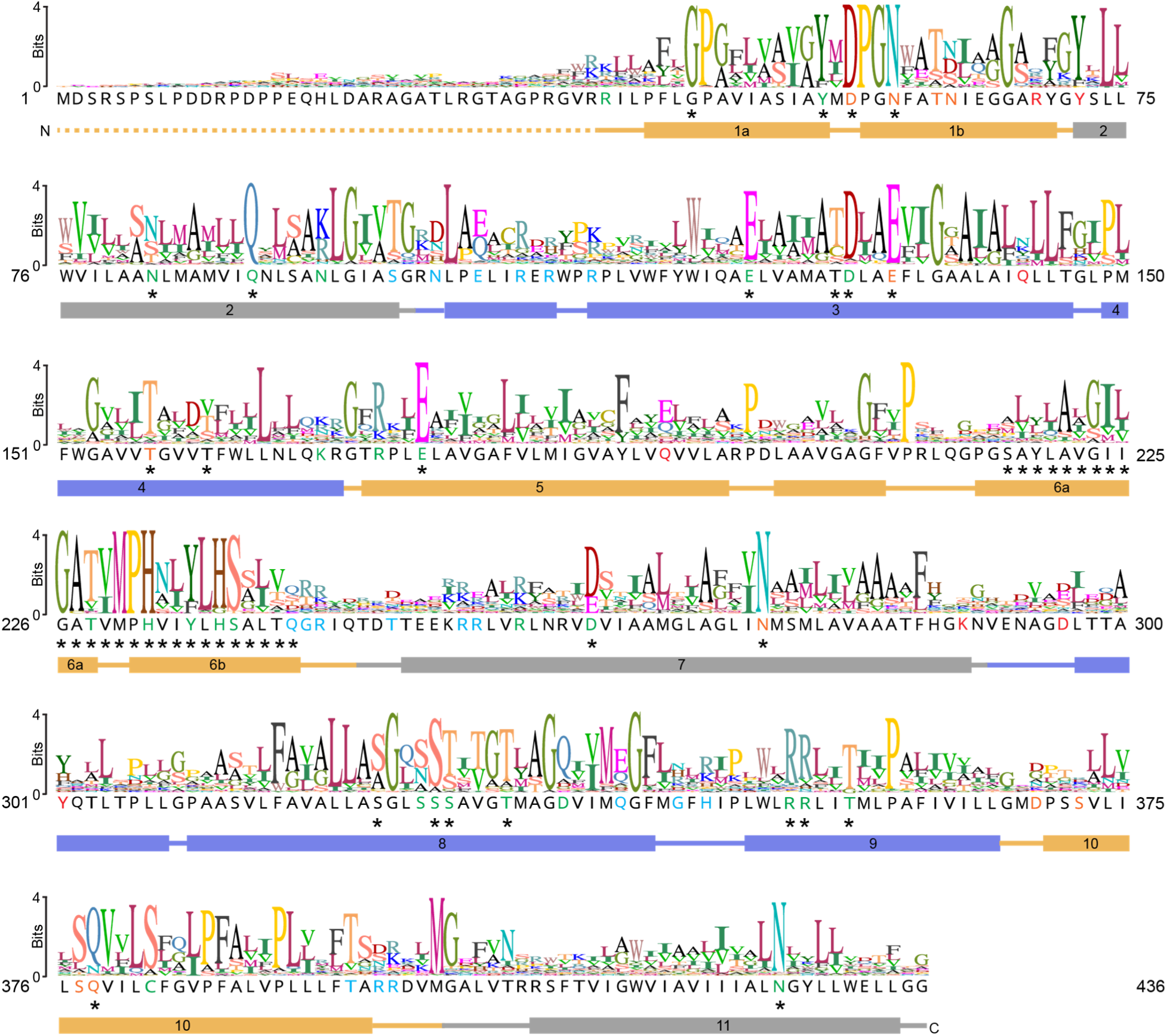
Secondary structure of DraNramp and sequence logo from Nramp alignment. The primary sequence of DraNramp is annotated with the secondary structure seen in PDB # 6BU5 and 6C3I. The sequence logo was generated using Geneious version 9.1 (Biomatters) from an alignment of 6878 sequences containing the canonical Nramp TM1 “DPGN” and TM6 “MPH” motifs (19). Conserved hydrophilic positions (defined as Ser + Thr + Tyr + Asn + Gln +Asp + Glu + His + Lys + Arg > 80%) are colored in DraNramp’s sequence consistent with the location in the structure as depicted in Figure 1. * indicates residues for which the effect of mutagenesis on transport ability was determined.

**Figure S2.**
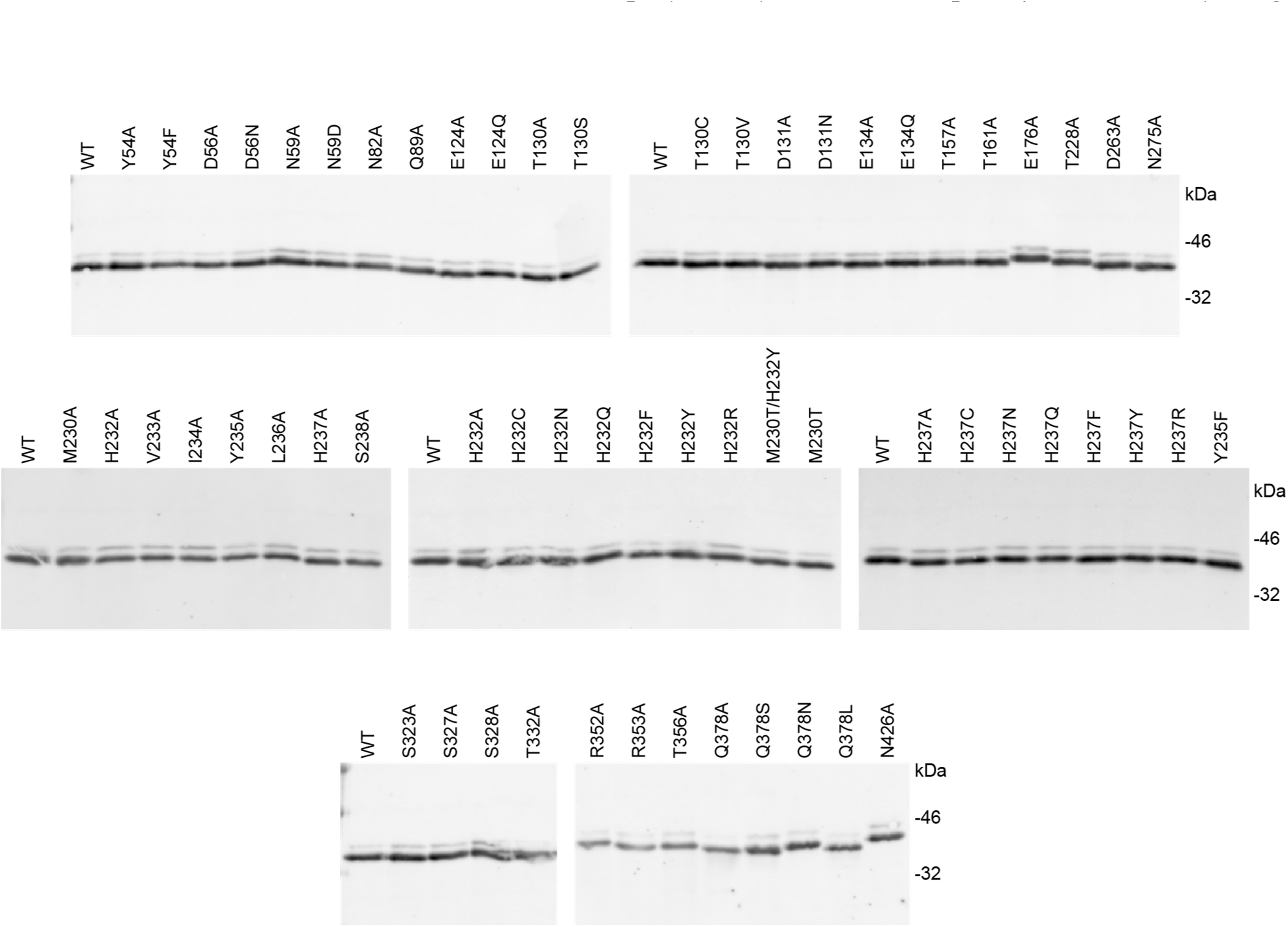
Expression of DraNramp mutants in C41(DE3) *E. coli*. Western blots for the N-terminal 8xHis-tag show that all Nramp point mutants discussed in this paper expressed in *E. coli*, most at a level similar to WT.

**Figure S3.**
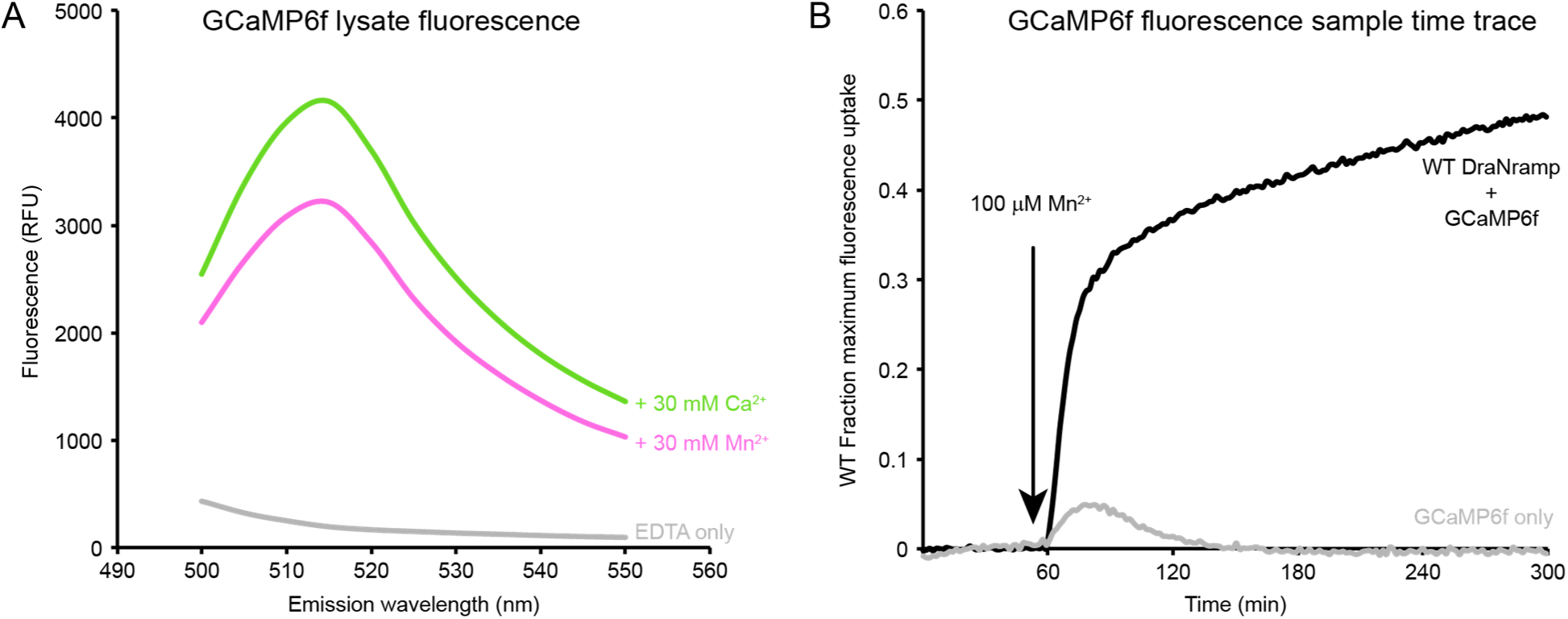
Validation of GCaMP6f as a Mn2+-sensitive fluorescent reporter. (A) C41(DE3) cells expressing GCaMP6f were lysed, EDTA was added to sequester native Ca^2+^, the lysate was buffered to pH 7.3, and excess additional Ca^2+^ (green) or Mn^2+^ (pink) was added before measuring fluorescence at an excitation wavelength of 470 nm. The gray trace corresponds to no added metal. (B) Sample time traces comparing relative fluorescence increases observed for C41(DE3) cells co-expressing WT DraNramp and GCaMP6f (black) and expressing GCaMP6f only (EV control; grey). The maximum fluorescence was determined by adding 10 mM Ca^2+^ to separate aliquots of cells.

